# Characterization of two novel EF-hand proteins identifies a clade of putative Ca^2+^-binding protein specific to the Ambulacraria

**DOI:** 10.1101/2020.05.22.110411

**Authors:** Arisnel Soto-Acabá, Pablo A. Ortíz-Pineda, José E. García-Arrarás

## Abstract

In recent years, transcriptomic databases have become one of the main sources for protein discovery. In our studies of nervous system and digestive tract regeneration in echinoderms, we have identified several transcripts that have attracted our attention. One of these molecules corresponds to a previously unidentified transcript (*Orpin*) from the sea cucumber *Holothuria glaberrima* that appeared to be upregulated during intestinal regeneration. We have now identified a second highly similar sequence and analyzed the predicted proteins using bioinformatics tools. Both sequences have EF-hand motifs characteristic of calcium-binding proteins (CaBPs) and N-terminal signal peptides. Sequence comparison analyses such as multiple sequence alignments and phylogenetic analyses only showed significant similarity to sequences from other echinoderms or from hemichordates. Semi-quantitative RT-PCR analyses revealed that transcripts from these sequences are expressed in various tissues including muscle, haemal system, gonads, and mesentery. However, contrary to previous reports, there was no significant differential expression in regenerating tissues. Nonetheless, the identification of unique features in the predicted proteins suggests that these might comprise a novel subfamily of EF-hand containing proteins specific to the Ambulacraria clade.

## Introduction

Modern genome and transcriptome studies allow for the identification and discovery of hitherto unknown sequences that code for different types of proteins. This discovery process has been possible due to the ease by which DNA and/or RNA sequences are obtained, even from non-model organisms that make available millions of sequences for comparative analyses. Our group has focused on transcriptomes obtained from normal and regenerating tissues of an echinoderm, the sea cucumber *Holothuria glaberrima* [1–4]. Studies in this model have been done to explore gene expression of intestinal and nervous systems in an attempt to expand our knowledge of the Echinodermata, a phylum which lies on the evolutionary branch of chordates [5,6].

In this effort, we have constructed several transcriptomic libraries using high throughput sequence analyses, including EST (expressed sequence tag) analyses [3], 454 and Illumina sequencing [7,8]. Moreover, we have performed differential gene expression studies, particularly microarrays and transcriptomic comparisons between normal and regenerating tissues. The results from these experiments have been a large number of differentially expressed genes associated with the regenerating tissues.

Out of these hundreds of genes, we have focused on the study of unknown sequences that show increased expression during regeneration. One of these molecules corresponds to a previously unidentified transcript from *H. glaberrima* that was shown to be upregulated during the initial stages of intestine regeneration by microarray analyses [3,8]. The sequence was annotated to public databases as *Orpin* (GU191018.1, ACZ73832.1) on 12-13-2009.

We now provide a full report on the putative Orpin sequence including the prediction of an N-terminal signal peptide, which is characteristic of secreted proteins. Moreover, we have discovered an additional *Orpin* isoform in *H. glaberrima* and provide a full description of both *Orpin* isoforms. Both sequences are newly discovered putative EF-hand coding proteins with structural characteristics that are evolutionarily related to this group of proteins. We have also probed other available databases and have found previously undescribed sequences whose similarities suggest they are part of the Orpin family, a protein family that appears to be restricted to the Ambulacraria clade.

## Materials and methods

### Ethics statement

This research deals only with invertebrate animals, thus the University of Puerto Rico IACUC waives ethical approval of research performed on invertebrates. Animals were sacrificed by immersion in ice cold water for 29-30 min and then sectioning the anterior part of the animal close to the oral nerve ring, which accounts for the main component of the nervous system.

### Animals

Adult specimens (10–15 cm in length) of the sea cucumber *H. glaberrima* were collected in coastal areas of northeastern Puerto Rico and kept in indoor in aerated seawater aquaria at room temperature (RT: 22°C ± 2°C). Evisceration was induced by 0.35 M KCl injections (3–5 mL) into the coelomic cavity [1]. Eviscerated animals were let to regenerate for 3, 5, 7, 10, and 14 days before the dissection and tissue extraction. For the dissection, organisms were anesthetized by placement in ice-cold water for 1 h. [1,3,9]. Dissected tissues were rinsed in ice-cold filtered seawater and processed for RNA isolation.

### RNA extraction and cDNA synthesis

RNA extraction was performed on tissue extracts of normal and 3, 5, 7, 10, and 14 dpe animals. Extracted tissues included gonads, mesentery, haemal system, respiratory tree, longitudinal muscle, and radial nerve cords. After dissection, tissues were placed in 1 mL of TRIzol reagent (Invitrogen), homogenized with a PowerGen Model 125 Homogenizer (Thermo Scientific) and incubated 30 min on ice. These samples were mixed vigorously with 200 μL of chloroform and incubated 10 min at RT. After centrifuged at 12,000 rpm at 4°C, the aqueous RNA phase was separated, mixed with 70% ethanol, and transferred to an RNeasy Mini Kit column (QIAGEN) for deoxyribonuclease (DNase) treatment (QIAGEN). Total RNA was extracted following the manufacturer’s protocol. The concentration and purity of the total RNA was measured using a NanoDrop ND-1000 spectrophotometer (Thermo Scientific). The cDNA was synthesized from 1 μg of the total RNA using the ImProm-II Reverse Transcription System (Promega) and oligo (dT)23 primers.

### Semi-quantitative RT-PCR

RT-PCR reactions were performed using cDNAs prepared from extracted RNA. These reactions were set up in a reaction volume of 25 μL with the final concentration of the PCR primers of 100 nM. Specific primers for the most variable regions between *Orpin A* and *Orpin B* sequences were designed using OligoAnalizer tools from the Integrated DNA Technology webpage (www.idtdna.com). The primers used were: *Orpin B* forward: 5’-ACAGGGAGTACAAACAGTCGTCAA-3’ and *Orpin B* reverse: 5’-CTATTTACTCTGCAACTGACACTTTCT-3’; *Orpin A* forward: 5’-ACTTCTGCAGAATCAGTTGTTAAGA-3’ and *Orpin A* reverse: 5’-TTCAGTGGAGTCGCCAAC-3’. RT-PCR reactions were performed on three independent RNA samples purified from each of the regeneration stages (previously mentioned) as well as from the normal intestines. The PCR amplification was done by an initial denaturation step of 94°C (45 s), a primer annealing step of 50.2°C (45 s), and an extension step of 72°C (45 s) with a final additional 72°C (10 min) for 28 cycles for *Orpin A*, 26 cycles for *Orpin B*, and 26 cycles for *NADH*, as the amplification parameters for each pair of primers. All samples were analyzed in triplicate. Additional tissues were amplified for 35 cycles (2–4 replicates). The relative expression of *Orpin A* and *Orpin B* was normalized relative to the expression of the housekeeping gene *NADH dehydrogenase subunit 5* using ImageJ software [10] from the optical density values from electrophoresed sample bands on 1% agarose gels, using a Molecular Imager ChemiDoc XRS+ (BioRad). The primers used for the *NADH* sequence amplification were: forward: 5’-CGGCTACTTCTGCGTTCTTC-3’ and reverse: 5’-ATAGGCGCTGTCTCACTGGT-3’. The *Orpin A* and *Orpin B* sequences were confirmed by sequencing excised electrophoresed sample bands at the Sequencing and Genotyping Facility (UPR-RP).

### Bioinformatics analyses

Homolog sequences were identified and retrieved from the NCBI GeneBank protein database [11] using the original Orpin sequence previously identified [8] as a query. BLASTp [12,13] were performed against the public non-redundant protein database in GeneBank. Conserved domain identification and UTR analysis were performed using CDD [14], RegRNA [15], UTRScan [16] and PSIPRED [17,18], ScanProsite [19], InterProScan 5 [20,21], Phobos [22], SignalP 5.0 [23], and Phobius [24] on Geneious 11.1.5 software (https://www.geneious.com). Sequence alignments were carried out with MUSCLE [25] (10 iterations) and the Blosum62 matrix and edited with Geneious software 11.1.5 (https://www.geneious.com). Note: It is possible that there are N-terminal sequencing artifacts on two annotated sequences from *A. japonicus* sequences (ARI48335.1 and PIK49419.1). If we delete the residues from the predicted cytoplasmic N-terminal region from the ARI48335.1 sequence and from PIK49419.1 up until their next methionine, they also show a predicted signal peptide of 21 residues each.

### Phylogenetic analysis

EF-Hand proteins and other similar sequences were retrieved from literature and protein database as mentioned in results section and the multiple sequence alignment was performed on MAFFT v7.309 [26] with BLOSUM62 scoring matrix, gap open penalty of 1.57, and offset value of 0.123. For the tree building, the Maximum-Likelihood analysis was done using JTT model of sequence evolution with 1000 bootstraps using PhyML 3.0 [27] plugin using Geneious 11.1.5 software (https://www.geneious.com). The corresponding sequences are included in S1 Table. The tree was edited for better visualization and colors in iTOL v4 online tool [28]. The (frog) *X. laevis*, (mouse), *M. musculus*, and (human) *H. sapiens* calcineurin A sequences were selected as outgroups and does not contain EF-Hand motifs. In addition, the Orpin homologs from *A. japonicus* ARI48335.1 and PIK49419.1 were edited for the analyses by deleting the N-terminal residues down to the second predicted methionine for the reason mentioned above.

### Statistical analyses

Statistical significance of the resulting data was evaluated through one-way ANOVA using the JMP^®^, Version 12. SAS Institute Inc., Cary, NC, 1989-2019. The multiple comparison procedure and statistical test Tukey-Kramer HSD (honestly significant difference) was used to determine significant differences between means from optical densities determined by ImageJ software as mentioned before [10]. The Tukey-Kramer results are displayed as small circles for high number of data points and large circles for low number of data points. The large red circle shows significant differences to small grey circles sample means. All values were reported as the mean ± standard mean error, including mean diamond with confidence interval ([1 − alpha] x 100), and outlier box plot from a quantiles report. While a *P* < .05 and *P* < .001 were considered to indicate statistical significance difference between groups.

## Results

### Identification of the original *Orpin* (*Orpin A*) sequence and characterization of a second *Orpin* isoform (*Orpin B*)

The original report [3] described a contig sequence (4766-1) which was later annotated as *Orpin*. This contig was used as a template to identify the remaining nucleotides upstream from the open reading frame (ORF) region through RACE-PCR analysis [29]. The *Orpin* sequence is composed of 106 nucleotides from the 5’ UTR and 291 nucleotides from the 3’ UTR with a 366 nucleotide ORF (plus stop codon) that encodes a putative 122 amino acid peptide followed by a stop codon (Figs 1 and 2). The nucleotide composition of this gene sequence was validated by sequencing the RT-PCR products amplified from a normal intestine tissue cDNA sample (Fig 2). At the time it was annotated in the NCBI database (ACZ73832.1; 12/13/2009), there was no match with other sequences. Two similar sequences from the hemichordate *Saccoglossus kowalevskii* were later added as *Orpin*-like sequences (XP_006824981.1 and XP_002736736.1).

**Fig 1.**
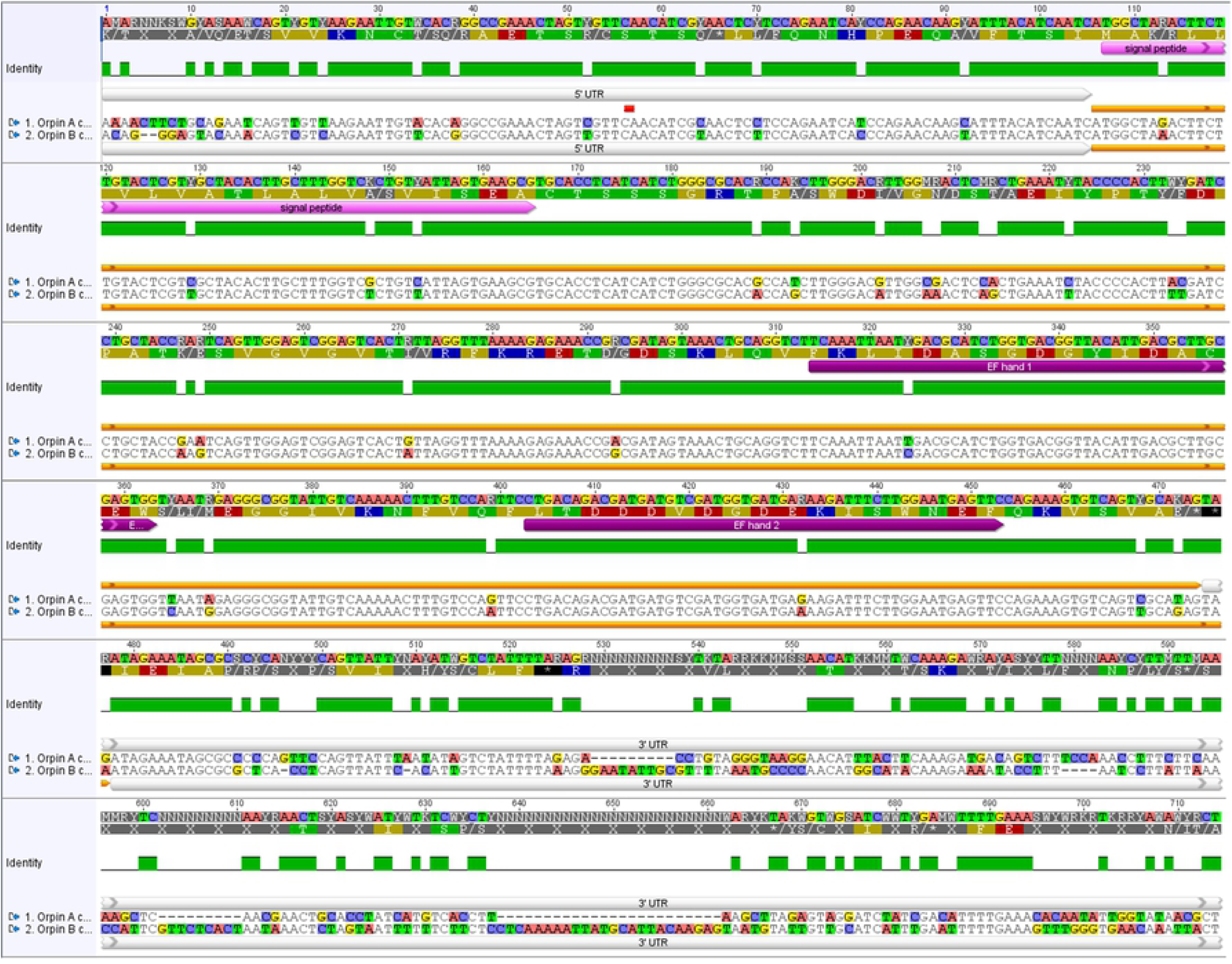
*Orpin A* and *Orpin B* are isoforms. Differences between sequences are highlighted. White bars: 5’ UTR and 3’ UTR regions of both sequences; pink bar: predicted signal peptides; orange bar: ORF regions; purple bars: predicted EF-hand motifs; green: conservation level; top sequences: nucleotide and amino acid consensus sequences. Differences between nucleotide sequences are highlighted. It is shown a significant difference, especially between both 3’ UTR sequences. Analysis was done using the MAFFT plugin in Geneious 11.1.5.

**Fig 2.**
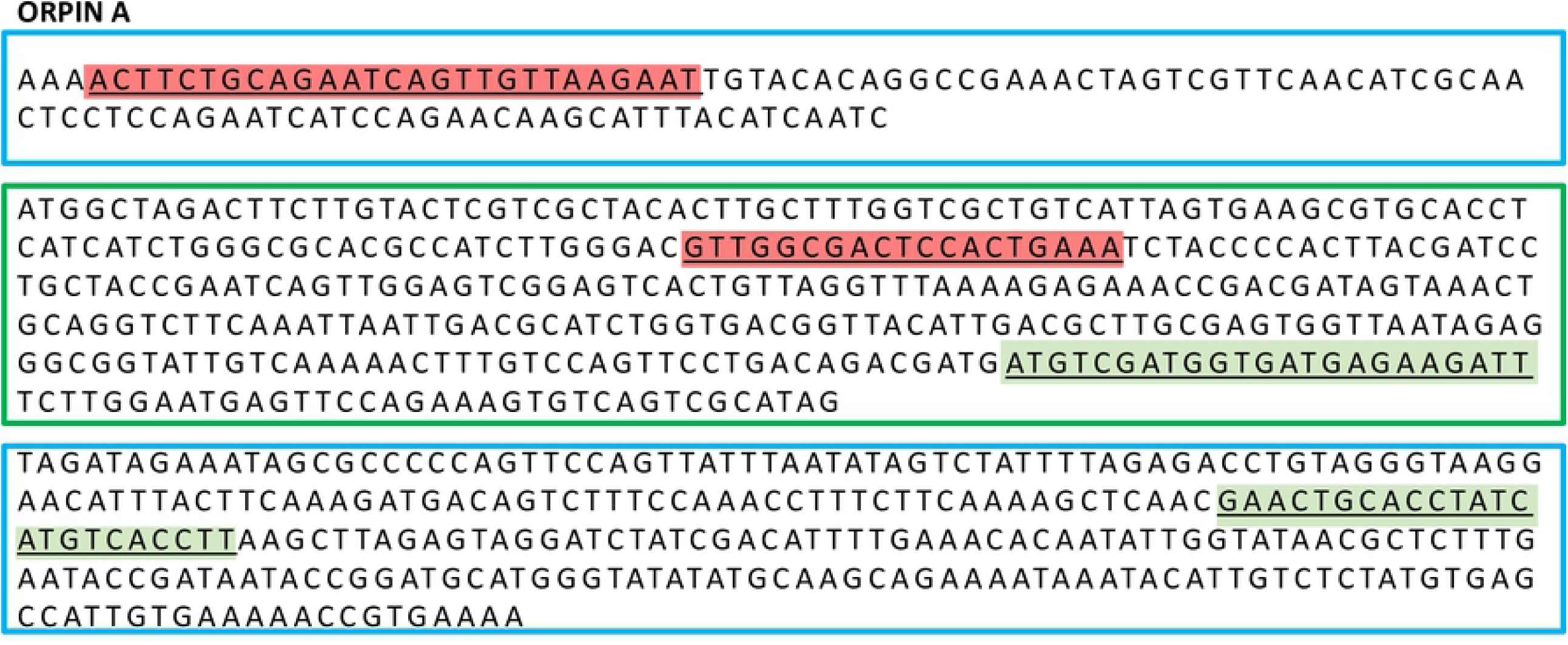
Primers for sq-RT-PCR of *Orpin A.* *Orpin A* UTRs regions (blue boxes) and coding region (green box) of the *Orpin A* gene. Primer sequences designed to specifically amplify *Orpin A* (light red letters). Primer sequences used for the identification of the original *Orpin* sequence in previous reports (green letters). These primers were designed prior to identification *Orpin* isoform.

After performing further in-depth analyses of the available transcriptome libraries from regenerating and non-regenerating intestine and regenerating and non-regenerating radial nerve, we discovered an additional highly similar sequence that was identified as a putative *Orpin* isoform. This new putative protein shared 90% identity and 98% similarity with the original *Orpin* sequence but displayed different UTR’s from the original sequence. We refer to this sequence as *Orpin B* to differentiate it from the original *Orpin* which we refer from now on as *Orpin A*.

The sequence corresponding to *Orpin B* was also validated through RT-PCR amplification and sequencing (Fig 3 and S1 Fig). *Orpin B* mRNA sequence is composed of 103 nucleotides from the 5’ UTR and 364 nucleotides from the 3’ UTR (Figs 1 and 3). Its ORF is 369 nucleotides (plus stop codon) long and encodes a putative 123 amino acid protein.

**Fig 3.**
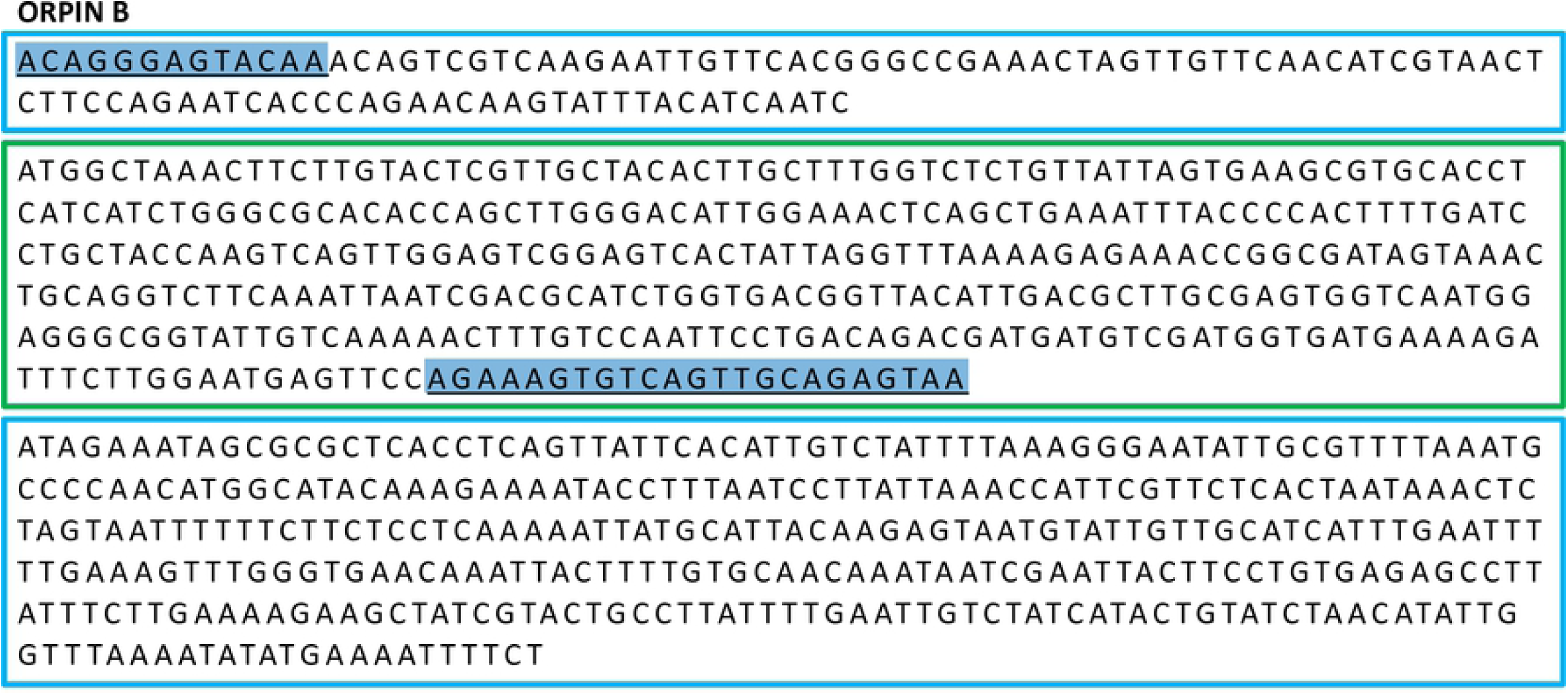
Primers for sq-RT-PCR of *Orpin B.* *Orpin B* UTR sequences (blue boxes) and coding region (green box). Primer sequences designed to specifically amplify *Orpin B* (blue letters).

Sequence comparisons among the two *Orpins* from *H. glaberrima* and the two *Orpin*-like sequences from *S. kowalevskii* show that the latter shared 46–50% identity and 76–77% similarity with the *Orpin A* (Fig 4). Similarly, *Orpin B* translated amino acid sequence shared 46–50% identity and 66–67% similarity with the sequences from *S. kowalevskii* (Fig 4). Furthermore, we identified three additional putative *Orpin* homologs from another sea cucumber species, *Apostichopus japonicus*, one from the starfish *Acanthaster planci*, and two from the sea urchin *Strongylocentrotus purpuratus* with expected values (*E*-value < 0.001 and total scores > 47.8). All *Orpin*-like sequences contain one domain that is predicted to be a calcium-binding domain composed of two EF-hand motifs at their carboxy-terminal (Figs 5 and 6).

**Fig 4.**
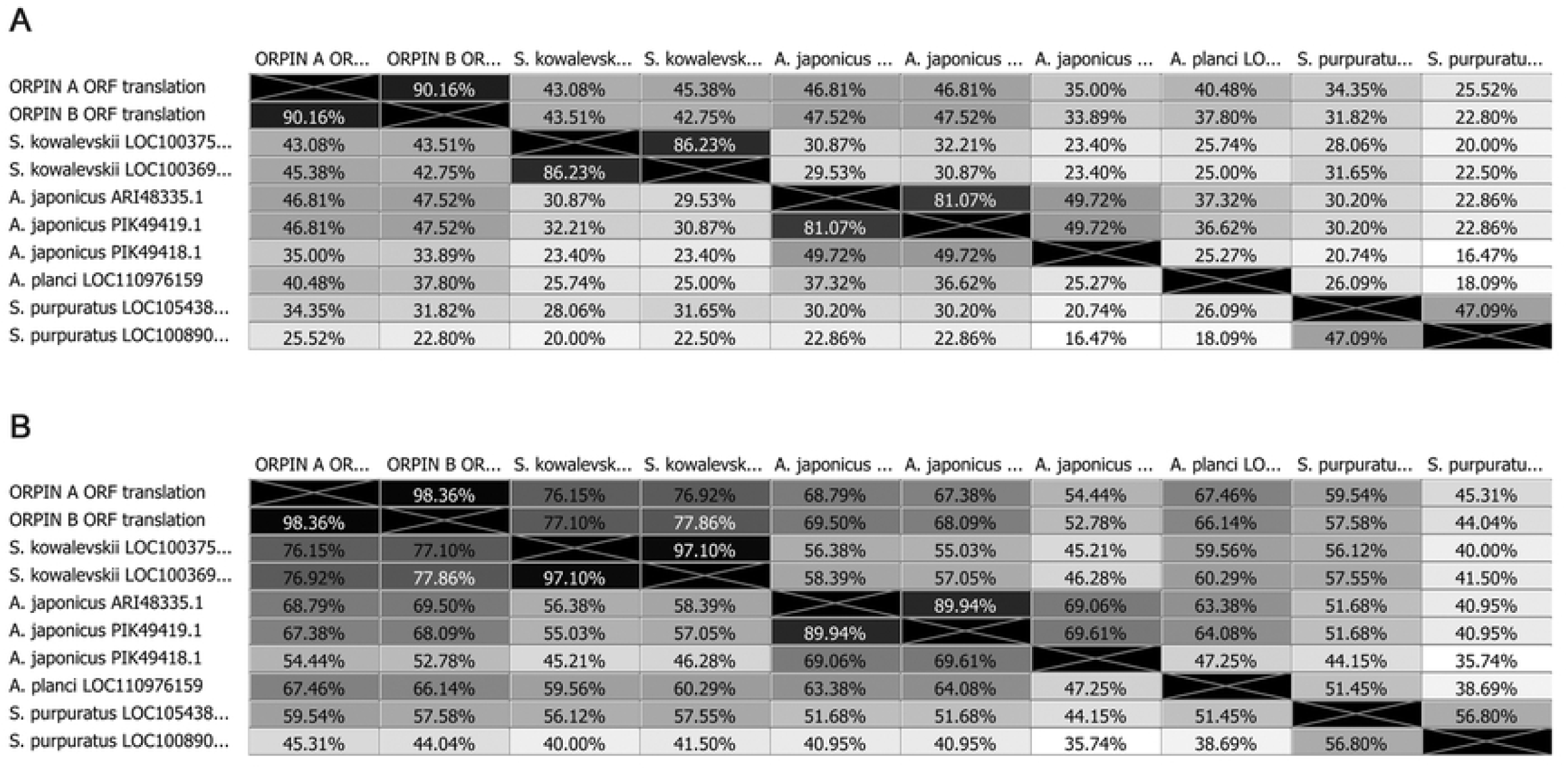
Orpin homologs pairwise sequence divergence. Translated amino acid sequences comparison by (A) identity% and (B) similarity%. The alignments were done using Muscle with 50 iterations using Geneious 11.1.5.

**Fig 5.**
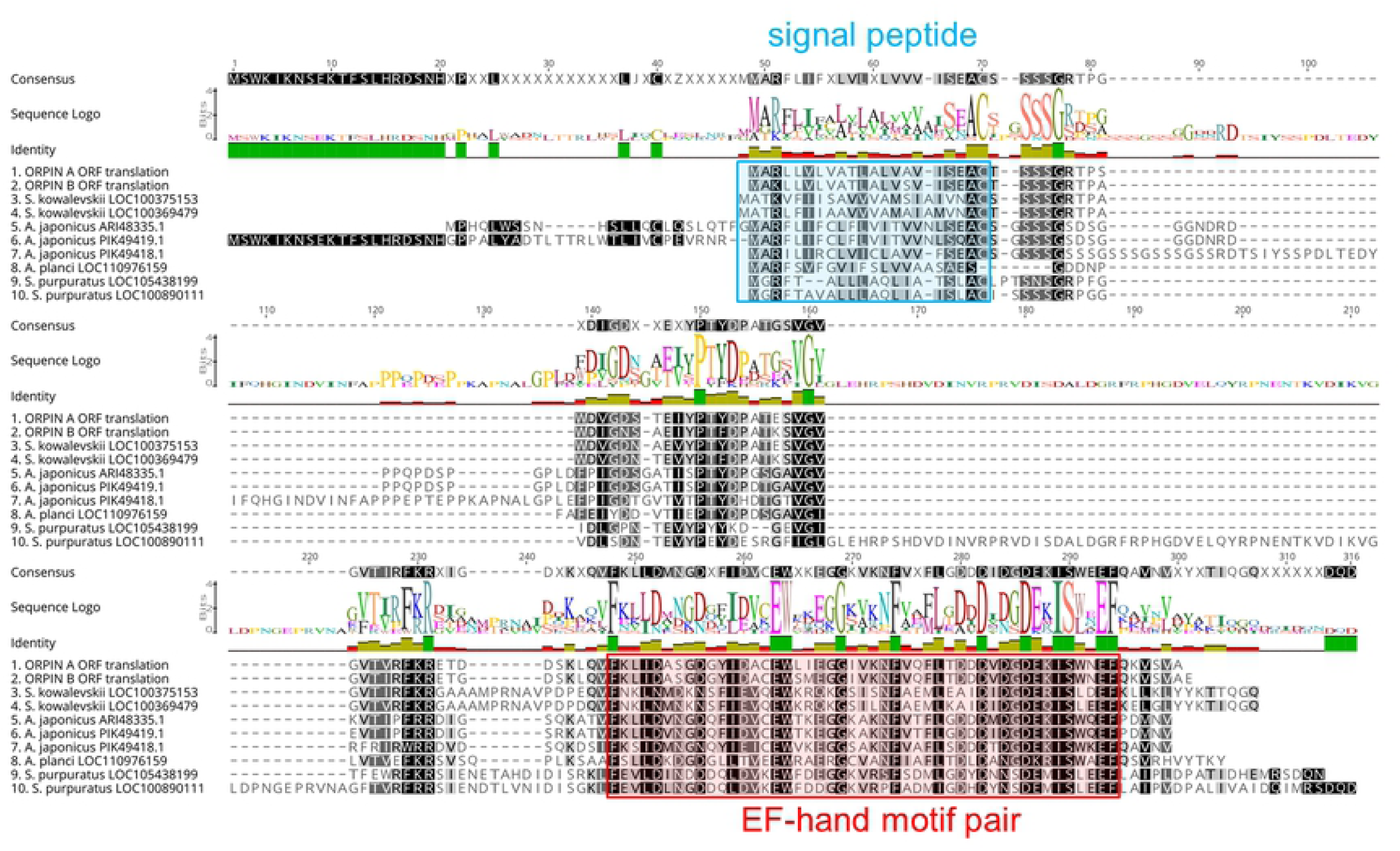
Orpin homologs alignment. The most conserved residues are indicated by letters in black boxes, green identity regions, and large cartoon letters at the sequence Logo. The exception is PIK49419.1 because 20 amino acid residues from the N-terminal portion are not compared to other sequences. We can see the additional N-terminal regions from *A. japonicus* sequences ARI48335.1 and PIK49419.1 that did not match to the other homologs. Blue box: signal peptide prediction; red box: EF-Hand motif pair prediction. This alignment was done by Muscle plugin with 50 iterations using Geneious 11.1.5.

**Fig 6.**
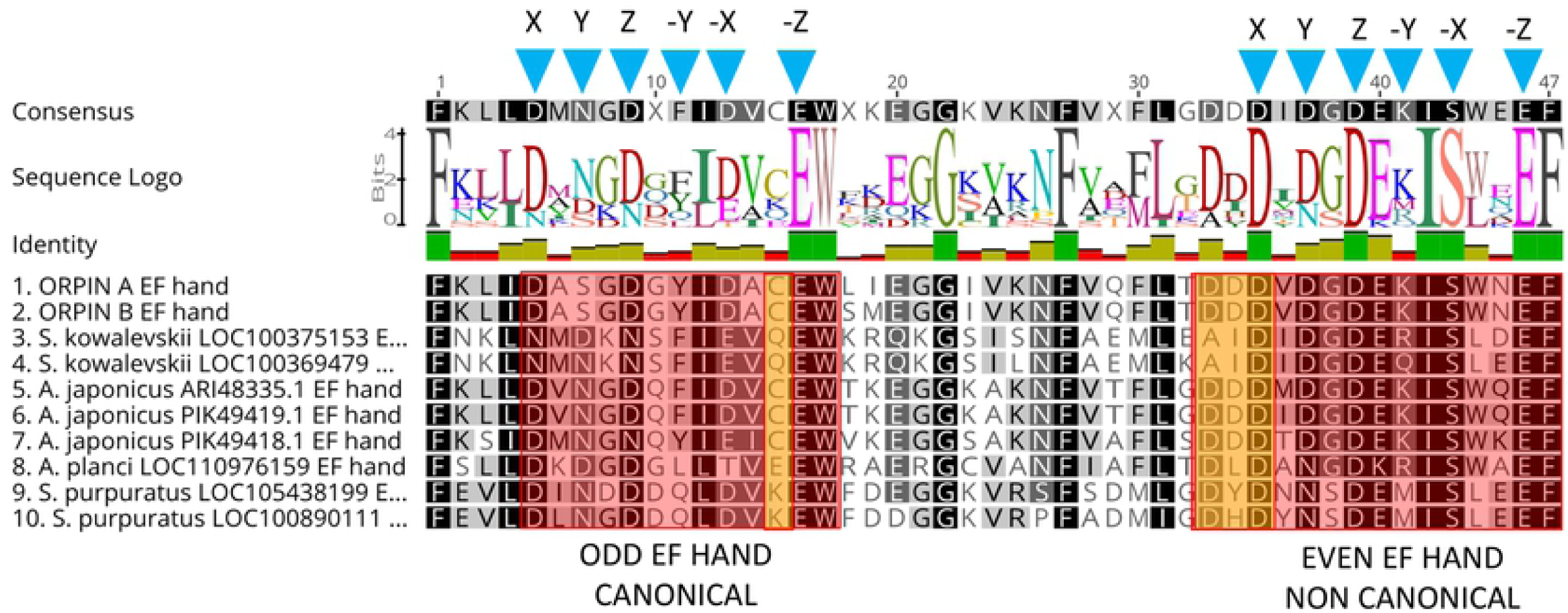
Orpin homologs EF-hand motifs alignment. The predicted odd EF-hands match with the canonical EF-hand pattern and the predicted even EF-hands were identified as non-canonical motifs (14 residues vs 12 residues) (red boxes). The non-canonical motifs are similar to vertebrates S100s. Predicted calcium coordinating residues from the identified EF-hands patterns are indicated by blue triangles. Holothurian Orpins contain a Cys residue at the −Z−1 position of the predicted calcium-binding loop (left orange box). The characteristic residues from Orpin residues are highlighted by orange boxes. Alignment was done using the MAFFT plugin in Geneious 11.1.5.

### Domain analyses

Orpin A and Orpin B amino acid sequences were analyzed using different bioinformatics tools (refer to methodology) for evidence that could point towards a possible function. After evaluating these sequences for domain composition using InterProScan and NCBI’s CDD, and Phobos [14,20–22], we identified that both sequences contain putative calcium-binding domain regions. Both Orpin isoforms shared identical calcium-binding loops residue composition. The key residue positions that participate in calcium chelation within these loops are conserved when compared with other known EF-hand proteins. The X, Y, Z, −X, −Z positions from each loop of the EF-hands are Asp, Asp, Asp, Asp, Glu (“odd loop”) and Asp, Asp, Asp, Ser, Glu (“even loop”), respectively (Fig 6). The only difference is located downstream to the “odd” loop. There are two consecutive amino acids, Leu87 and Ile88, immediately after the Trp86 (−Z+1) of the first calcium-binding loop from Orpin A which are changed to Ser87 and Met88 in Orpin B. Interestingly, Orpin A and Orpin B included a Cys residue at −Z−1 position which is particular to both sea cucumber sequence homologs and is an unusual feature in EF-hand proteins.

When we compared *H. glaberrima* Orpin EF-hand sequences to the other identified putative homologs from *S. kowalevskii, A. japonicus, A. planci*, and *S. purpuratus*, we found that additional positions are highly conserved as well. All Orpin sequences share conserved positions at X−4 (Phe), −Z (Glu), −Z+1 (Trp) and −Z+6 (Gly) positions from the “odd” EF-hands, and X−8 (Phe), X (Asp), Z (Asp), −X−1 (Ile), −X (Ser), −Z (Glu) and −Z+1 (Phe) positions from the “even” EF-hands. Alternatively, there are residues particular to the EF-hands from *H. glaberrima* Orpin isoforms, such as Ala at X+1, Ser at Y, Ala at −X+1 from the odd EF-hand, and Val at X+1 and Asn at −X+2 from the even EF-hand. Moreover, there are residues that are particular to the holothurians such as Lys at X−3, Cys at −Z−1, Lys at −Z+9 from the odd EF-hand, and Lys at −Y from the even EF-hand (Fig 6).

Additional bioinformatics analyses revealed the presence of a signal peptide in the N-terminal of both isoforms (Fig 7). These signal peptides are 20 amino acids long each and are mainly composed of hydrophobic residues. The predicted signal peptides of Orpin A and Orpin B are nearly identical, with the exception of two residues at positions 3 (Arg/Lys) and 15 (Ala/Ser). Furthermore, InterProScan and Phobius identified the same region of 20 residues as a possible transmembrane region. In both cases, a high probability of a cleavage site was identified at the Cys21 residue of each isoform. If the signal peptide is eliminated, the remaining sequence is predicted to be localized outside the cytoplasm (Fig 7). This strongly suggests that these peptides could be secreted to the extracellular space and not targeted to the membrane of other cell organelles.

**Fig 7.**
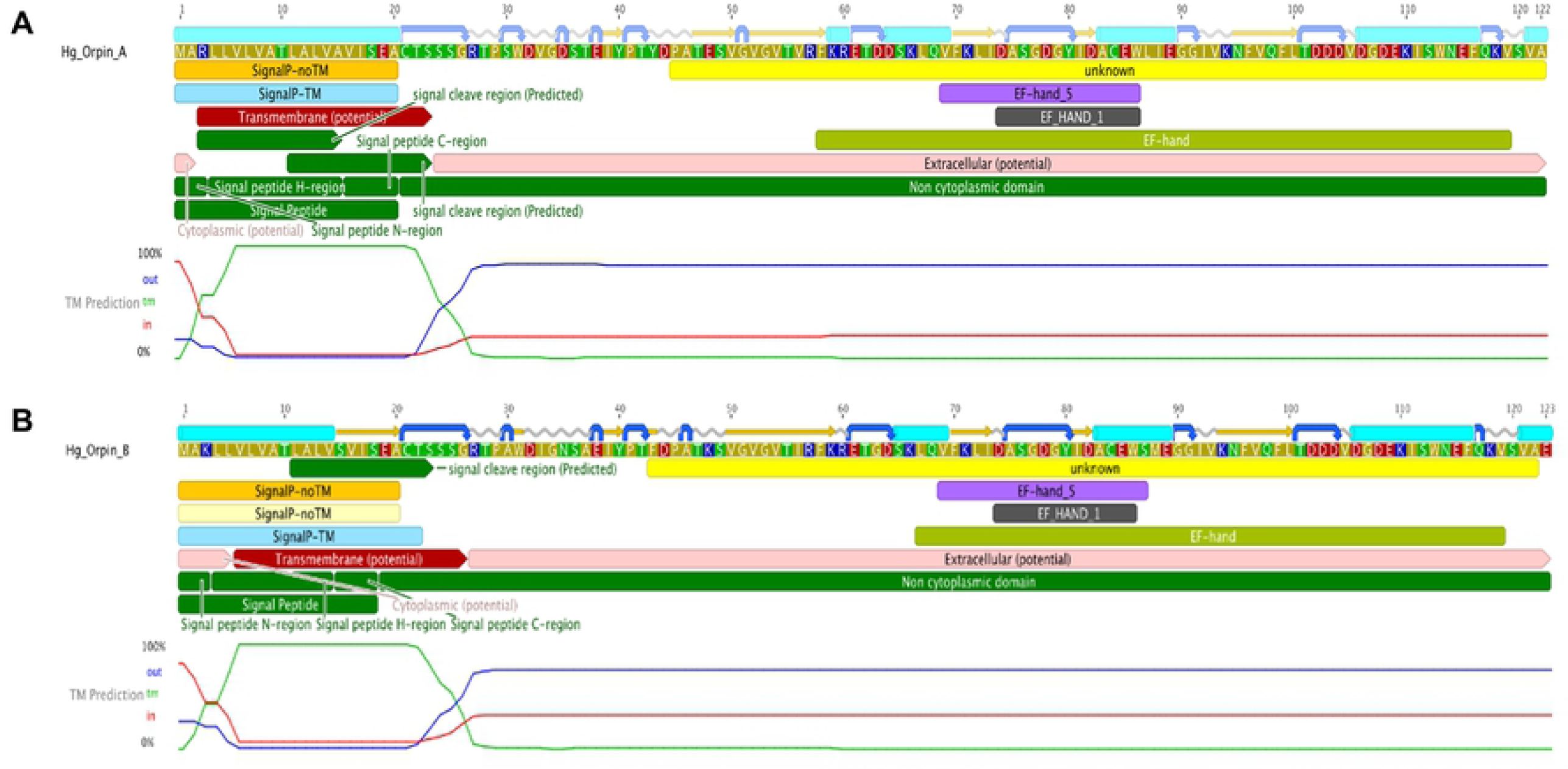
Orpin A and Orpin B bioinformatics characterization. (A) Orpin A and (B) Orpin B have predicted signal peptides at their transmembrane N-terminal regions including cleavage sites. Also, the two isoforms have predicted EF-hand motifs in their non-cytoplasmic regions. These were predicted by various bioinformatics plugin tools using Geneious 11.1.5.

The average length for the predicted signal peptides of the Orpin homologs is 20–22 residues based on SignalP and Phobius predictions [23,24,30], from the initial Met residue. The predicted signal peptides from the two *S. kowalevskii* sequences are longer (22 residues) than the other Orpins. The predicted signal sequence of a sea urchin homolog (XP_011664021.1) is the shortest (18) of the Orpins (Fig 5).

The UTR’s of the *H. glaberrima Orpin* sequences were also analyzed. Even though the 5’ UTR’s from *Orpin A* and *Orpin B* are 80.2% identical, there are 20 nucleotide differences between them, mainly SNPs. A ribosome binding site within the 5’ UTR of each isoform sequence was identified. In contrast, the retrieved 3’ UTR’s of both *Orpin* isoforms were completely different. Polyadenylation sites were identified in both *Orpin A* [8] and *Orpin B* downstream to their corresponding stop codons. Interestingly, these analyses revealed the presence of two putative Musashi binding elements (MBEs) within the 3’UTR of *Orpin A*. Even though the available retrieved 3’ UTR from *Orpin B* is longer than its paralog, no MBEs were identified within this sequence. Surprisingly, two putative MBEs were also found within the coding sequence of each *Orpin* isotype.

### Orpin phylogenetic analysis

In order to determine the relationship of the different Orpin homologs among themselves and with other EF-hand proteins, a phylogenetic tree was constructed with the PhyML program using a MAFFT alignment as input [26,27]. Orpin A and Orpin B amino acid sequences were used as probes to identify the closest sequences through BLAST searches against the public databases. In addition, representative sequences from different EF-Hand subfamilies of various organisms were obtained from the scientific literature and available databanks. These sequences included members from the following protein families: S100s, calcineurin, recoverin, calbindin, parvalbumin, oncomodulin, osteonectin, SPARC, troponin C, calmodulin, centrin, Spec, and recoverin (S1 Table). Thus, these sequences were used for the final alignment to generate the phylogenetic tree.

The results from this analysis cluster Orpin and the identified hypothetical homologs from *S. kowalevskii* (acorn worm), *A. japonicus* (sea cucumber), *A. planci* (sea star), and *S. purpuratus* (sea urchin) together with a bootstrap value of 92, separately from other subfamilies of EF-Hand proteins (Fig 8). The other EF-Hand protein sequences cluster together as individual groups. The Orpin-like cluster was the most distant group after the outgroup sequences of mouse and frog calcineurin A, which do not contain EF-Hand motifs, suggesting that Orpins have evolved separately and are not direct homologs of EF-Hand proteins from other species. As expected, *H. glaberrima* Orpins were close to the other sea cucumber *A. japonicus* Orpin-like sequences. The most distant Orpin homologs were those from sea urchin *S. purpuratus.* The closest protein cluster was the osteonectins, BM-40, or SPARC proteins, which comprise a group of secreted CaBP modulators with a single pair of EF-Hand motifs. After these, the other group of proteins that appeared close by were the S100s, which also are small secreted proteins with two EF-hand motifs. The tree also showed the other outgroup EF-hand lacking protein, calcineurin A from humans, was placed separately from the other EF-Hand proteins of a high number of motifs (3 to 6 EF-Hands).

**Fig 8.**
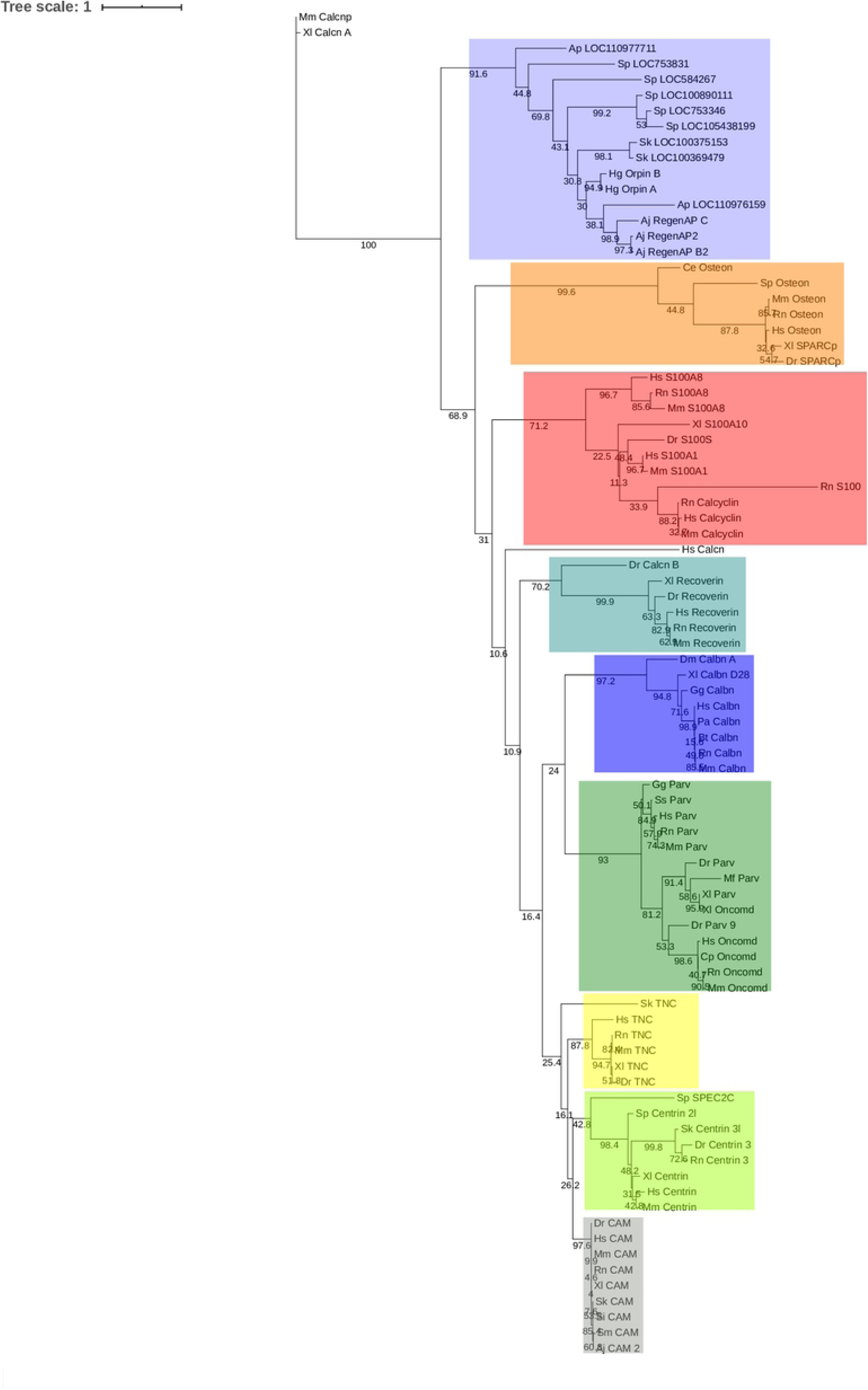
Orpin isoforms are specific to the Ambulacraria clade. EF-hand protein representative sequences from different subfamilies were aligned to build a phylogenetic tree. Orpin homologs were clustered together as a group, separated to the other EF-hand proteins. The tree was made using the PhyML plugin ran through Geneious 11.1.5. The parameters used for this analysis were JTT model of amino acid substitution and 1000 bootstraps. Scale bar: 1. Protein sequences accession numbers are included in S1 Table.

### *Orpin* gene is expressed in several tissues of *H. glaberrima*

In order to determine the distribution of *Orpin* expression, mRNA was obtained from different tissues or organs of normal (non-regenerating) *H. glaberrima* specimens and processed for PCR analysis. The tissues and organs selected were: small intestine, large intestine, mesentery, radial nerve complex, longitudinal body wall muscle, gonads, and respiratory tree. Primers were designed for the specific detection of *Orpin A* and *Orpin B* mRNA sequences (Figs 2 and 3). Transcript levels were evaluated relative to the expression of *NADH subunit 5*, a constitutively expressed housekeeping gene. The results showed that *Orpin A* and *Orpin B* shared similar tissue specificity (Figs 9 and 10). Transcripts were detected in the gonads, muscle, mesentery, and haemal system but not in the respiratory tree nor in the nerve. Tissue expression varies significantly, with higher expression levels in the mesentery followed by the expression in muscle and gonads where it is slightly higher than in other tissues. Interestingly, a faint second lighter band was detected from *Orpin B* samples from gonads and muscle tissues.

**Fig 9.**
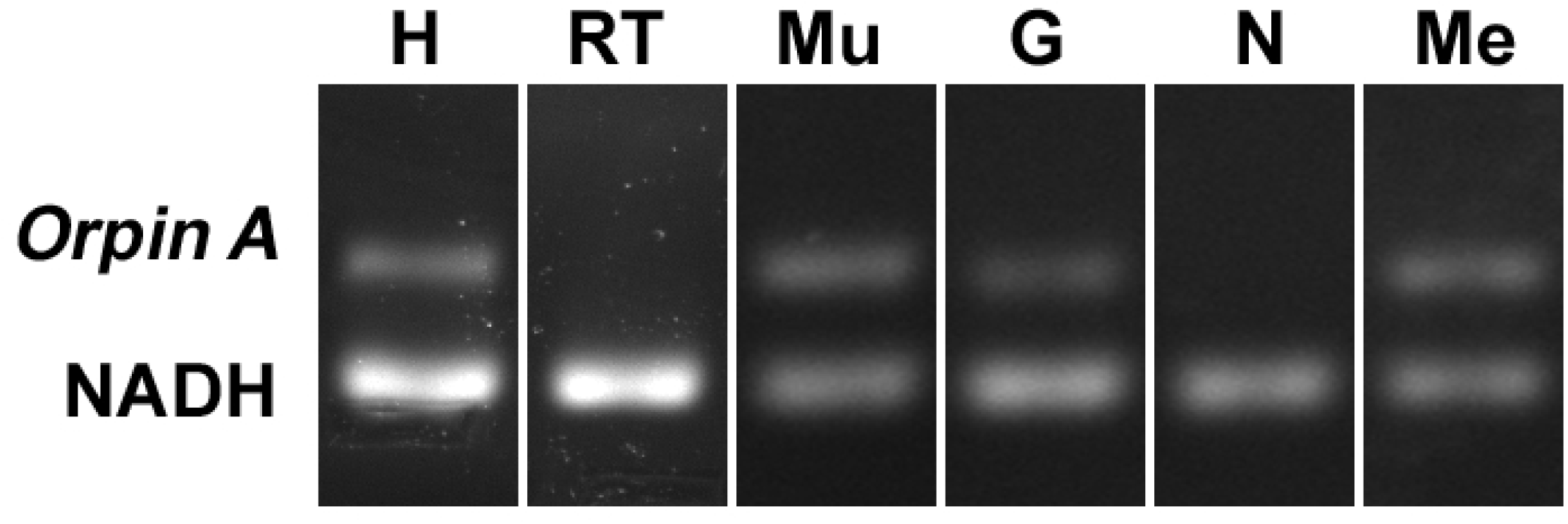
*Orpin A* expression in different tissues. Composite image from RT PCR amplification of *Orpin A* from *H. glaberrima* tissues. *Orpin A* expression (top band) was detected in haemal system (H), muscle (Mu), gonads (G), and mesentery (Me) relative to the expression of *NADH. Orpin A* was detected neither in the nerve (N) nor in the respiratory tree (RT). The image is a composite from different gels and is divided by a white line.

**Fig 10.**
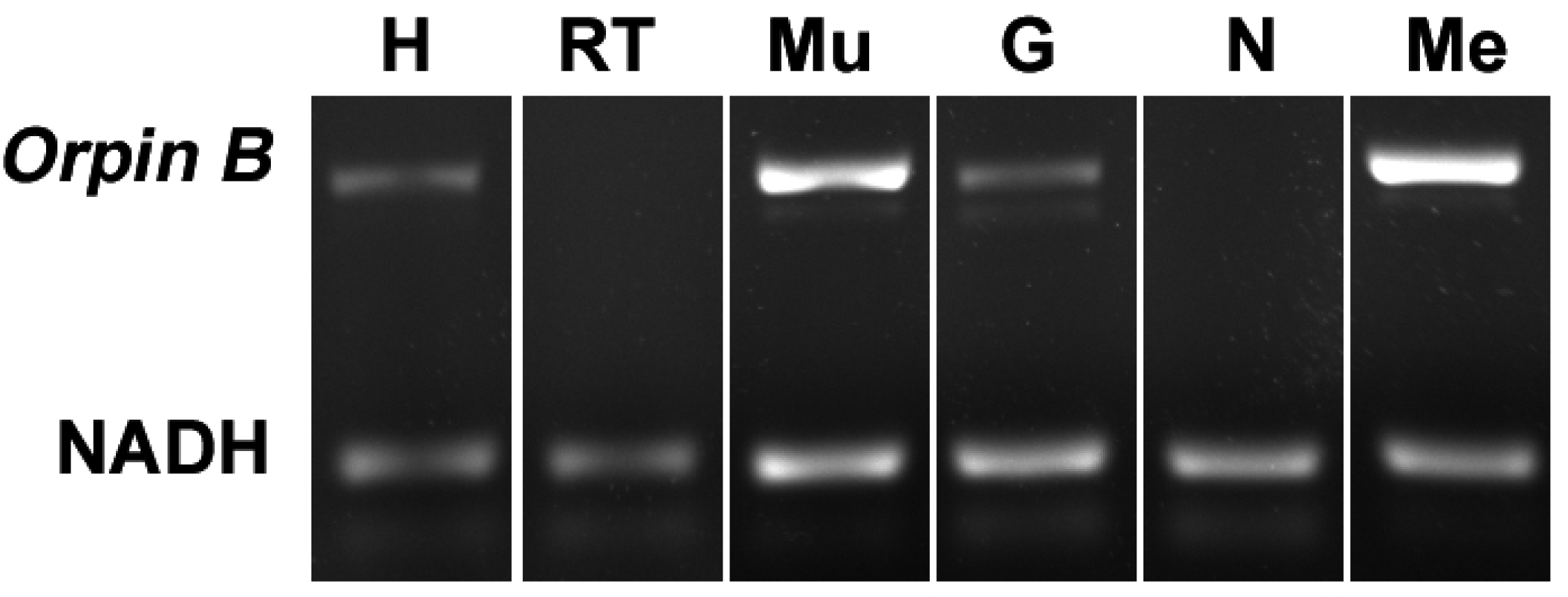
*Orpin B* expression in different tissues. Composite image from RT PCR amplification of *Orpin B* from *H. glaberrima* tissues. *Orpin B* expression (top band) was detected in haemal system (H), muscle (Mu), gonads (G), and mesentery (Me) relative to expression of *NADH. Orpin B* was detected neither in the nerve (N) nor in the respiratory tree (RT). The faint band below the *NADH* band corresponded to primer dimers. The image is a composite from different gels and is divided by a white line.

### *Orpin* expression during intestinal regeneration in the sea cucumber *H. glaberrima*

Previous results from our laboratory have shown that *Orpin* was differentially expressed in regenerating intestinal tissues when compared to normal intestinal tissues [3,8]. In order to validate the upregulation of this novel sequence during regenerative processes, *Orpin* transcript levels were measured during different stages of intestine regeneration. In contrast to previous experiments where no particular effort was made to separate the intestine of normal animals from the attached mesenteries, in the present experiments we measured separately the intestine (a mixed portion from the small intestine and from the large intestine) and the mesentery that attaches the intestine to the body wall, for the normal (non-regenerating) samples. Orpin transcript levels were measured relative to the housekeeping gene *NADH subunit 5*. The gene expression levels were monitored using semi-quantitative RT-PCR of tissue extracts from 3 days post evisceration (dpe), 5 dpe, 7 dpe, and 10 dpe along with tissues from normal intestine and normal mesentery.

Previously it was found that the expression levels of *Orpin A* increased after 3 days of intestine regeneration and then gradually returned to the basal levels at 14 days of regeneration. In contrast, to those findings [3,8], there was no statistically detected difference found between the transcript expression of *Orpin A* from normal intestine samples and those from any of the studied regenerative days (Data not shown). This was also true for *Orpin B* (Data not shown). However a high differential expression was detected between tissues from *Orpin A* from normal mesentery and tissues from 7–10 dpe sample group, with a *P* < .05 (*P*=.002) (Fig 11). *Orpin B* exhibited a high differential expression between tissues from normal mesentery and tissues from 3–5 dpe with a *P* < 0.05 (*P*=.02), and 7–10 dpe with a *P* < .001 (Fig 12).

**Fig 11.**
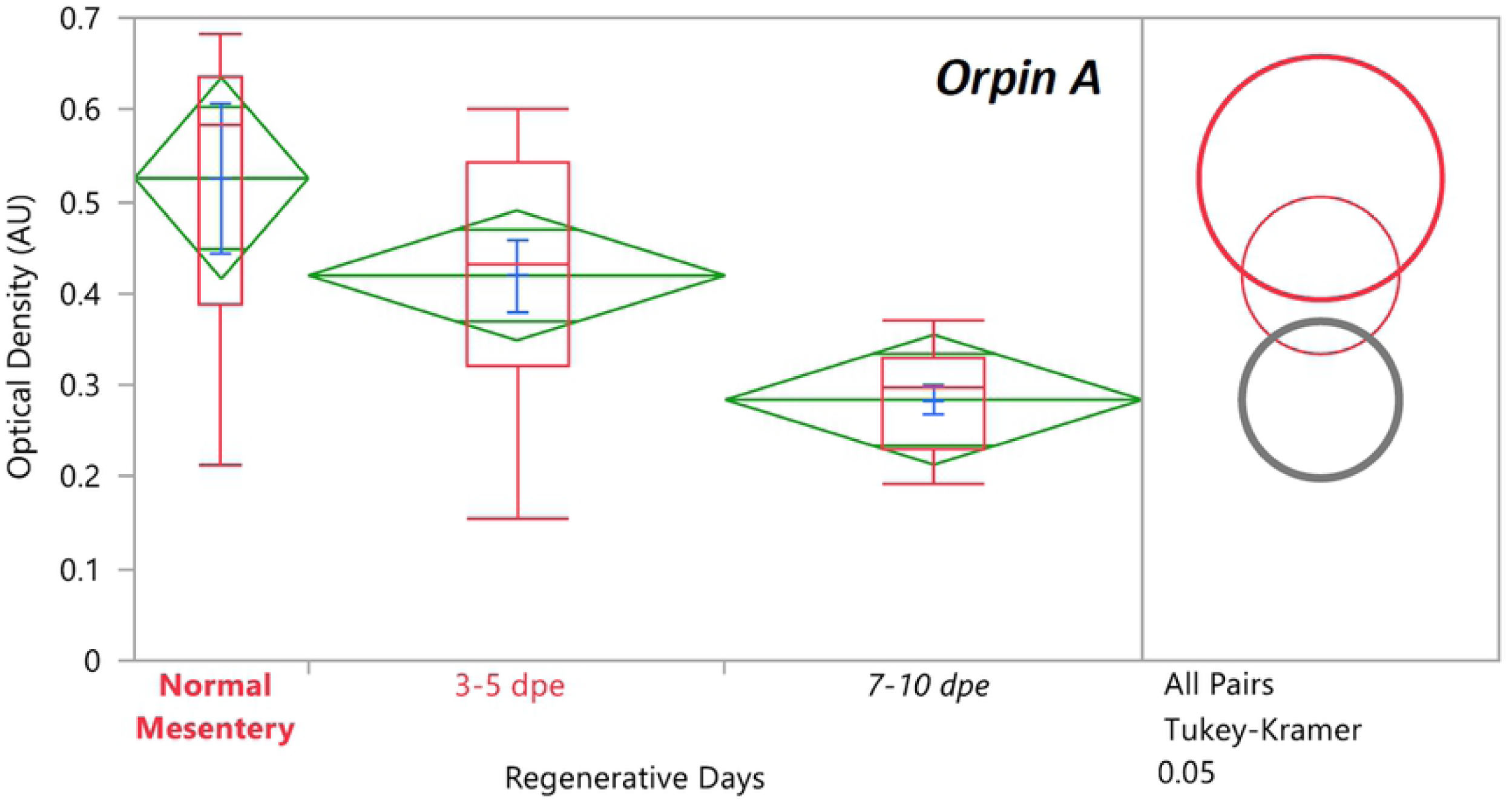
*Orpin A* expression during intestine regeneration grouped tissues. Semi-quantitative RT-PCR amplification of *Orpin A* transcripts from mRNA samples from different intestine regenerative days compared to the corresponding expression in samples from normal intestine (NI) and normal mesentery (NM). A statistical high differential expression was found between NM and 7–10 days post evisceration (dpe) (*P* < .05; *P* = .002); and between 3–5 dpe and 7–10 dpe (*P* < .05; *P* = .03) as indicated in the all pairs Tukey-Kramer HSD test. The large red circle (low number of data points) displays the significant difference between the small grey circles (high number of data points) group means. Red boxes: outlier box plots summarizing the distribution of points at each factor level from the quantiles report. Green diamonds: sample mean and confidence interval ([1 − alpha] x 100). Blue lines: standard mean error. JMP^®^, Version 12 software was used for the statistical analyses.

**Fig 12.**
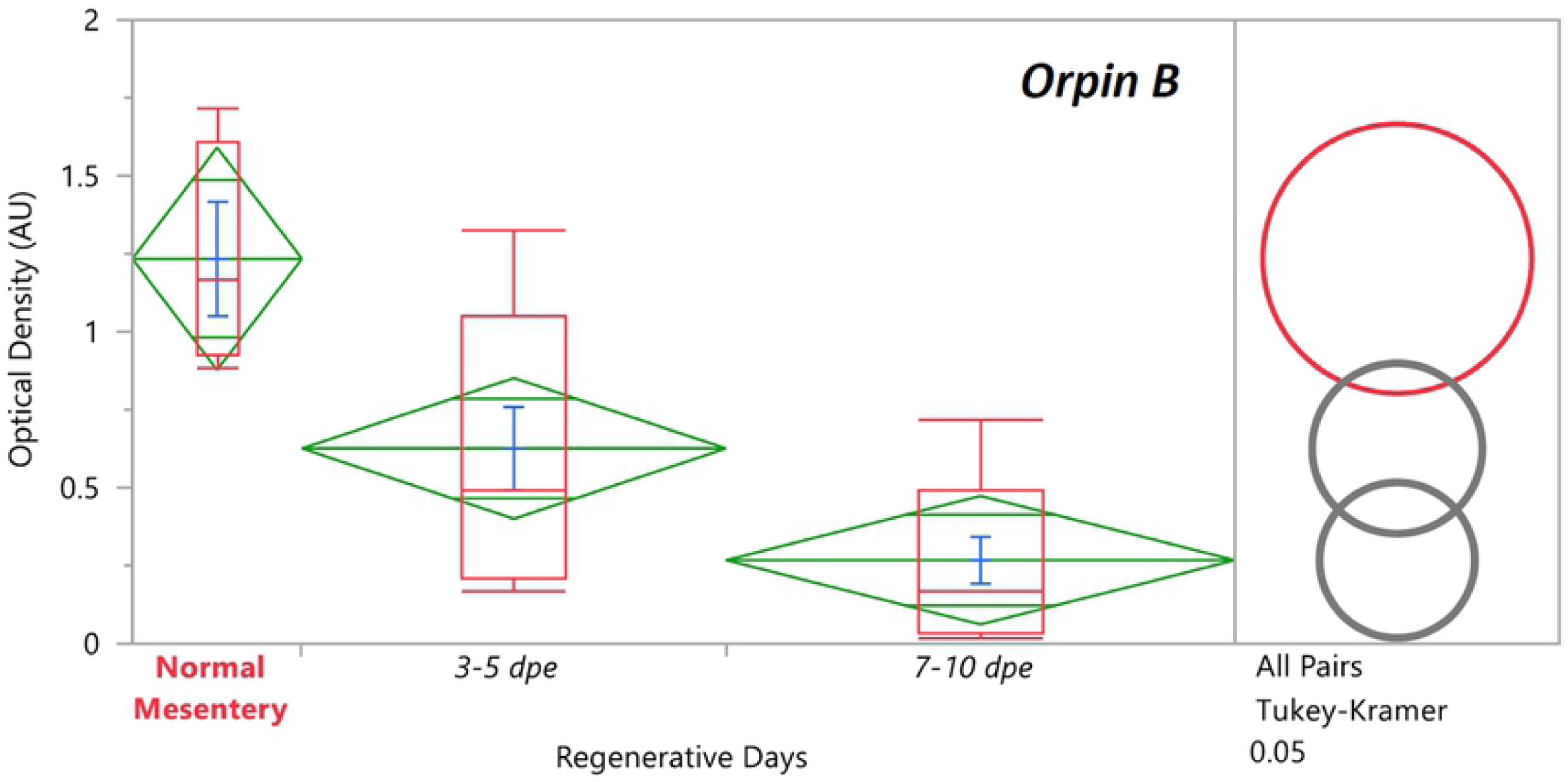
*Orpin B* expression during intestine regeneration grouped tissues. Semi-quantitative RT-PCR amplification of *Orpin B* transcripts from mRNA samples from different intestine regenerative days compared to the corresponding expression in samples from normal intestine (NI) and normal mesentery (NM). A statistical high differential expression was found between NM and 3–5 dpe (*P* < .05; *P* = .02), and to 7–10 dpe (*P* < .001) as indicated in the all pairs Tukey-Kramer HSD test. The large red circle (low number of data points) displays the significant difference between the small grey circles (high number of data points) group means. Red boxes: outlier box plots summarizing the distribution of points at each factor level from the quantiles report. Green diamonds: sample mean and confidence interval ([1 − alpha] x 100). Blue lines: standard mean error. JMP^®^, Version 12 software was used for the statistical analyses.

Interestingly, we found a different expression profile between *Orpin A* and *Orpin B* transcript levels. While both *Orpin* forms show subsequently decreases in their expression to similar levels at 7-dpe to 10-dpe, the decrease of *Orpin B* seems to occur much faster than that of *Orpin A*.

## Discussion

We have now described the presence of two predicted EF-hand domain-containing proteins from the sea cucumber *H. glaberrima*. These putative proteins apparently belong to a unique group that is present in echinoderms and hemichordates. According to the mRNA distribution in the sea cucumber, the translated proteins are expressed in multiple organs. Moreover, they are highly represented within the mesentery of the normal and regenerating intestine. The possibility that these are Ca^2+^-binding proteins is discussed in the following sections.

### *Orpins* are novel genes

When the first *Orpin* sequence from *H. glaberrima* was identified, no other sequence that showed significant similarity to it could be found within databases [3,8]. A few months later, two highly similar sequences (and possible homologs) were identified in the hemichordate *Saccoglossus kowalevskii* (acorn worm) and were added to the databases. These sequences were annotated with accession numbers XM_006824918.1, XP_006824981 (*E*-value: 9E-31) and XM_002736690.2, XP_002736736 (*E*-value: 2E-29). Later on, several homologs from closely related organisms of the Echinodermata phylum were added to the public databases: two from the sea urchin *Strongylocentrotus purpuratus*, three from other sea cucumber *Apostichopus japonicus*, and one from the starfish *Acanthaster planci*. The finding of an additional *Orpin* sequence in *H. glaberrima* increased to ten the known sequences and suggested that these sequences belong to a novel family of proteins within a group of metazoans. In all cases, the sequences have been annotated with little or no descriptive information other than their tissue/organism from where they originated. At present, *Orpins* appear to be restricted to the Ambulacraria clade (the group that encompasses echinoderms and hemichordates), however, it remains to be seen if, with the sequencing of other animal genomes, the specificity of *Orpins* to the Ambulacraria still stands.

Although the *H. glaberrima* A and B *Orpin* variants share a high percentage of similarity at the nucleotide and protein levels, our data suggest that they correspond to distinct genes. First, the nucleotide and putative amino acid differences are distributed throughout the complete sequence of both variants. Second, even though the 5’ UTR sequences of both *Orpin* sequences share a large similarity, they are not identical, and similar to the coding region, have multiple nucleotide differences distributed along the nucleotide sequences. Third, their 3’ UTR sequences are very different, sharing minimal similarities. Finally, other species also have more than one *Orpin*-like gene. For example, sequence information from *Orpin* homologs from *S. kowalevskii* and *S. purpuratus* were annotated as located in different loci. These differences are characteristic of different genes rather than allele variants or products from differential splicing. Nonetheless, in spite of these results that suggest two different isoforms originating from two different genes, it is necessary to have the genome information as conclusive evidence. Moreover, it should be emphasized that while *Orpin A* and *B* mRNAs have been identified and sequenced from various tissues (see below), the *S. kowalevskii, A. japonicus, A. planci*, and *S. purpuratus* sequences are hypothetical mRNA/protein-coding sequences that remain to be characterized. Even though it was expected that *H. glaberrima Orpin* sequences would be more similar to those from the other sea cucumber *A. japonicus*, the results showed that they shared higher similarity to the hemichordate *S. kowalevskii* sequences. This can be attributed to longer N-terminal regions from two of the *A. japonicus* sequences (ARI48335.1 and PIK49419.1) that did not match with any of the other homologs. These additional regions were annotated as part of the corresponding ORFs due to an identified methionine upstream to the one that matched the other homologs. Given the fact that these sequences were not validated, it has to be considered the possibility that these first encoding methionine residues could be the result of a PCR artifact, and could be in fact part of their corresponding 5’ UTR regions. Hopefully, the characterization of *Orpin* isoforms will provide essential insights that eventually would make feasible the characterization of these homolog sequences.

### Are Orpins calcium-binding proteins?

Rigorous analysis of the residue composition of the EF-hand domains coupled to structural and functional experimentation is the mainframe of the study of uncharacterized CaBPs. Thus, understanding the architecture of EF-hand domains provides a hint of the role that a particular EF-hand protein might have.

One of the most prominent characteristics of the putative protein Orpin is the presence of EF-hand motifs that are a distinctive signature of calcium-binding proteins. The EF-hand motif has been used as a standard of reference for the description of the calcium-binding loops from the corresponding domain by the residues at key positions for the chelation of each calcium ion. Even though the EF-hand motifs are a characteristic feature of many CaBPs, few of them consist of less than four EF-hands. As mentioned before, Orpin A and Orpin B paralogs comprise a single putative calcium-binding domain composed of two EF-hands. The canonical EF-hand motif topology is a helix-loop-helix conformation, which regularly binds calcium ions [31,32]. Usually, this conformation is composed of a highly conserved 12 residues calcium-binding loop flanked on both sides by alpha-helices [33]. The residues that participate in calcium coordination were labeled as X, Y, Z and −X, −Y, −Z [32,34], and those conserved positions are conventionally used as a reference frame to analyze the calcium-binding potential and dynamics of the EF-hand CaBPs. A typical EF-hand domain is composed of two calcium-binding loops containing motifs flanked by alpha-helices. The adjacent alpha helices are named incoming and exiting helices from the odd (N-terminal) and even (C-terminal) EF-hand motifs [35–38]. *H. glaberrima* Orpin isoforms odd calcium-binding loops are composed of highly conserved key amino acids of the canonical domain structure. We showed that these domains shared a high level of conservation in the residues that participate in the chelation of calcium ions. However, the “even” calcium-binding loop from sea cucumber Orpins slightly deviates from the canonical pattern. The conserved Gly6 residue that provides for loop flexibility is substituted by a Glu6. This substitution is well conserved throughout all the available Orpin homologs with the exception of the starfish sequence. Also, they share a conserved Trp13 at position −Z+1 of the “odd” EF-hand. Furthermore, this conserved residue seemed to be particular to Orpins after comparison to the other 84 EF-Hand sequences from this study. Interestingly, a Cys11 residue is particular to the holothurian Orpins.

In view of these facts, Orpin homologs can be classified as novel EF-hand proteins. Although the bioinformatics analysis strongly suggests that they might be a new type of CaBPs, in order to assure this, experimental confirmation of the actual binding of calcium ions will be required along with the phylogenetic evidence provided in this study.

### Orpin relationship with other Ca^2+^ binding proteins

Whether they are indeed CaBPs or not, the sequence comparisons show that Orpins share several characteristics with CaBP subfamilies and that in fact, CaBPs account for the most similar proteins in the database. Several patterns have been developed to accurately classify newly discovered EF-hand proteins. The main pattern is the calmodulin canonical EF motif mostly known as the DXDXDG pattern [39,40], which contrasts with the pseudo-EF-hand binding loops from the most recent vertebrate S100 family of proteins.

One of the main EF-hand protein subfamilies are the S100 proteins. These proteins are small CaBPs containing only two EF-hand motifs. Nevertheless, their N-terminal motifs are considered pseudo-EF-hands, which is the main characteristic of this protein family. At the moment of this study, this subfamily has only been found in vertebrates [41]. Although there were no pseudo-EF-hand predicted from both Orpin isoforms sequences, they share several characteristics with S100 proteins, such as the small size, acidic composition, secretion to extracellular location, and EF-hand motif number. Thus, Orpin sequences share structural features with the main CaBP subfamilies, making it difficult to classify them as any of them. Moreover, we have shown that Orpin and Orpin-like sequences clustered together more closely to osteonectins (BM-40/SPARC) proteins, a group of secreted CaBPs with only two EF-Hand motifs, suggesting that these two groups shared a common ancestral origin. In addition, the best-known CaBPs that grouped closely to Orpin-like sequences were mouse (*M. musculus*, AAA37432.1) and frog calcineurin A isoforms (*X. laevis*, AAC23449.1). These two sequences and the human calcineurin A (*H. sapiens*, AAC37581.1) do not contain EF-hands and were included as outgroups for this analysis. We emphasize the fact that the calcineurin isoform B from zebrafish does contain an EF-hand domain, thus it is separated from the outgroup calcineurin A sequences and is included with the other aligned CaBPs. These results suggest that Orpin sequences comprise a specific EF-hand protein group that is different from the other known subfamilies of calcium-binding proteins. These data suggest that we can be dealing with a new subfamily of EF-hand proteins that is specific to the Ambulacraria clade.

### Orpins as secreted proteins

In addition to their EF-hand motif, an additional feature of Orpins is the presence of a signal peptide. In this respect, Orpin isoforms strongly resemble the groups of EF-hand proteins that are secreted, namely the osteonectins, oncomodulins, and S100s. Similar to Orpin sequences, osteonectins (BM-40/SPARC), oncomodulin, and S100 proteins are small peptides containing two EF-hand motifs each and are secreted to the extracellular milieu. Interestingly, these proteins display calcium-mediated dimerization either as heterodimers as in the case of S100A8/S100A9 [42] or homodimers as in the case of S100P [43], osteonectins [44], and oncomodulin [45]. Usually, EF-hand proteins containing signal peptides are targeted to the outer plasma membrane. There, these CaBPs can act as growth factors recognizing binding targets located on other cell surfaces, thus activating different signaling pathways. Such is the case of osteonectin (BM-40/SPARC), which promotes changes in cell morphology, disrupt cell adhesion, inhibit cell cycle, regulate extracellular matrix, and modulate cell proliferation and migration [46], thus underlying the process of wound repair. Similarly, the secreted (although lacking a signal peptide) oncomodulin and S100 proteins, are involved in a variety of biological processes including: cell proliferation, differentiation, survival, nerve regeneration, interaction with transcription factors, and calcium homeostasis [47–55] among other functions.

### *Orpins* and regeneration

Previous results from our laboratory had shown differential expression of the *Orpin* transcript when mRNAs levels from 3- and 7-day regenerating intestines were compared to normal intestinal tissues. Thus, we had concluded that *Orpin* was over-expressed during intestinal regeneration. Our new data, where we detect high levels of the transcript in the mesentery region, questions our previous interpretation. Thus, if we consider that the samples containing the 3- and 5-day regenerating intestinal rudiments contain a large proportion of the remaining mesentery (that remains attached to the body wall), then the high expression of *Orpin A* and *B* transcript levels that were detected in the 3- or 5-day regenerating tissues can be interpreted as representing the expression in the mesenteric portion and not necessarily in the rudiment itself. As the rudiment itself grows and encompasses a larger proportion of the dissected tissues (in relation to the mesentery) then the *Orpin* expression would appear to decrease. This is why there is no difference between normal mesentery and early regenerating rudiments. The previously observed difference between regenerating rudiments and “normal” intestine would be merely a reflection of the proportion of mesenterial tissue present in both samples; low in “normal” intestines and high in regenerating ones.

In this respect, the isolated “normal” mesentery is a more appropriate control to compare the relative expression of *Orpin* transcript sequences between regenerating and normal tissues. This is particularly true in early regenerating stages (3–5 days) when the proportion of tissues corresponding to the mesentery is quite high.

While the lack of *Orpin* differential expression argues against a possible role in the intestinal regeneration process, we cannot completely exclude this possibility. The presence of Musashi binding elements within *Orpin* isoform sequences suggests that the post-transcriptional regulation of their mRNAs might be controlled by RNA-binding proteins. This element is present in genes that are post-transcriptionally regulated in a spatial and temporal dependent manner [56,57]. Moreover, this type of regulation has been implicated in the self-renewal of epithelial, neural and hematopoietic stem and progenitor cells [58–64]. Such is the case of the target transcript encoding the transcription factor TTK69 in *Drosophila*, where translational activation is mediated by the neural *Drosophila* Musashi. In this way, the Musashi protein induces the differentiation of *Drosophila* IIb cells as neural precursor cells by repressing the translation of the mRNA of this neural differentiation inhibitory factor [58]. Furthermore, the expression of mammalian Numb protein (m-Numb) induces the expression of regeneration-related genes such as prostate stem cell antigen (PSCA) and metallothionein-2 (Mt2) in gastric mucosal regeneration in mice. Musashi protein (Msi1) enhances the expression of m-Numb during this regenerative process through post-transcriptional regulation. Having stated this, we cannot disregard the possibility that *Orpin* isoforms play a role during the initial stages of intestine and nerve regeneration. Nonetheless, to explore this possibility, we need to determine if a Musashi protein is present in the *H. glaberrima* proteome during regenerative processes of the sea cucumber and that it binds to *Orpins* mRNA.

It is of interest that *Orpin* isoforms are found to be differentially expressed in a transcriptomic library of regenerating nerve from *H. glaberrima* [7], also suggesting a possible regeneration-associated function. The two sequences displayed an increase in expression during the regeneration of the radial nerve complex after induced injury. By day 2 and also by day 20 after nerve injury, *Orpin A* expression was significantly higher than non-regenerating radial nerve (*P* < .001). In addition, *Orpin B* was higher in the same samples after day 2, 12, and 20 after nerve injury (*P* < .001). However, in view of our findings in the intestinal system, it remains to be determined whether this differential expression is also the product of the gene is expressed preferentially in the remaining tissues following injury, and not necessarily of increasing its expression.

In summary, we have identified and characterized a group of Orpin-like proteins from a particular group of invertebrate deuterostomes and shown they all share similarities in size, domain composition, and little significant similarities to other known EF-hand protein sequences. We provide bioinformatics evidence for the presence of signal peptides and cleavage sites in these proteins that suggest secretion of the putative proteins to the extracellular environment. Together, with the identification of predicted EF-hand domains with unique features, we can suggest that these might comprise a novel subfamily of EF-hand containing proteins specific to the Ambulacraria clade. Finally, we studied their expression in normal and regenerating tissues, with the surprise finding that they are highly expressed in the intestinal mesentery.

## Acknowledgments

The authors thank Dr. Vladimir S. Mashanov for editorial comments on the manuscript as well as to kindly provide sequence differential expression information from *Holothuria glaberrima* regenerating nerve.

## Supporting information

**S1 Fig. RT-PCR primers test.** *Orpin A* and *Orpin B* RT-PCR amplification using primers for semi-quantitative analysis. cDNA used was from a pool of cDNAs including normal intestine, mesentery, and regenerating days tissues. Run cycles: 35. *Orpin B* primers show amplification of dimers, also shown in cDNA (-) control.

**S1 Table. EF-hand proteins sequences for the phylogenetic analyses**

